# Colocalization highlights genes in hypothalamic–pituitary–gonadal axis as potentially mediating polycystic ovary syndrome risk

**DOI:** 10.1101/2020.01.10.901116

**Authors:** Jenny C Censin, Jonas Bovijn, Michael V Holmes, Cecilia M Lindgren

## Abstract

Polycystic ovary syndrome (PCOS) is a common disease in women with consequences for reproductive, metabolic and psychological health. Women with PCOS have disrupted signalling in the hypothalamic-pituitary-gonadal axis and studies have indicated that the disease has a large genetic component. While a recent genome-wide association study of PCOS performed in up to 10,074 cases and 103,164 controls of European decent identified 14 PCOS-associated regions, much of the disease pathophysiology remains unclear.

Here, we use a Bayesian colocalization approach to highlight genes that may have a potential role in PCOS pathophysiology and thus are of particular interest for further functional follow-up. We evaluated the posterior probabilities of shared causal variants between PCOS genetic risk loci and intermediate cellular phenotypes in one protein and two expression quantitative trait locus datasets, respectively. Sample sizes ranged from 80 to 31,684. In total, we identified seven proteins or genes with evidence of a shared causal variant for almost a third of PCOS signals, including follicle stimulating hormone (FSH) and the genes *ERBB3*, *IKZF4*, *RPS26*, *SUOX*, *ZFP36L2*, and *C8orf49*. Several of these genes and proteins have been implicated in the hypothalamic-pituitary-gonadal signalling pathway.

In summary, our results suggest potential effector proteins and genes for PCOS association signals. This highlights genes for functional follow-up in order to demonstrate a causal role in PCOS pathophysiology.

## Introduction

Polycystic ovary syndrome (PCOS) is a common endocrinopathy, affecting between 6-10% of women of reproductive age (1). The disease has a heterogeneous clinical presentation (2–4), with consequences for reproductive, metabolic, and psychological health (2,3). Commonly, diagnosis is based on the Rotterdam criteria, which requires two out of three of 1) oligo- or anovulation, 2) signs of hyperandrogenism (clinical or biochemical), and 3) polycystic ovarian morphology, as well as exclusion of other diagnoses (3,4).

PCOS pathophysiology is still largely unclear (2), although one mechanism may be disrupted gonadotropin signalling that disturbs normal follicular development and ovulation (3). In healthy women of reproductive age, the pituitary gland secretes the gonadotropins luteinizing hormone (LH) and follicle-stimulating hormone (FSH) in response to pulsatile secretion of gonadotropin releasing hormone (GnRH) (5,6). These GnRH pulses are more frequent in women with PCOS (3,7). This changed secretion pattern causes an imbalance between LH and FSH, and a higher LH/FSH ratio (3,7–11), which may contribute to e.g. hyperandrogenism and disturbances in follicular maturation and ovulation (3,12). Other possible contributing factors that have been suggested include for example insulin resistance and inflammation (3,8). There is also evidence for a strong genetic component, with genetic factors suggested to explain 66% of the disease variance (13). Previous genome-wide association studies (GWAS) have highlighted risk loci close to genes with a plausible connection to PCOS pathophysiology, including genes involved in for example insulin and hypothalamic-pituitary-gonadal (HPG) signalling (e.g. *INSR*, the insulin receptor gene and *FSHR*, the FSH-receptor gene) (3,14–18). However, for most PCOS-associated loci the mediating genes and their functional effects remain to be identified and/or confirmed (17,18).

One approach to improve biological understanding of a disease risk locus is through colocalization analysis of the disease and intermediate cellular phenotypes, such as gene expression and protein levels in different tissues (19). Therefore, to improve understanding of PCOS pathophysiology, we investigated the evidence of colocalization between 14 PCOS-associated loci identified in a recent GWAS in Europeans (18) and one study with protein and two studies with expression quantitative trait loci (pQTL and eQTL, respectively). Our results highlight several genes and proteins linked to the HPG axis and follicular development, including e.g. FSH, *ZFP36L2*, and *RAD50*, that may be of particular interest for further functional follow-up.

## Results

### Colocalization highlights genes with a potential mediating role

We extracted 14 PCOS risk loci from a recent GWAS of up to 10,074 cases and 103,164 controls of European ancestry (Fig 1, Table 1) (18). We assessed the evidence for colocalization (19) between these loci and pQTL data from INTERVAL and eQTL data from the Genotype-Tissue Expression project (GTEx) and eQTLgen (20–23). PCOS summary statistics based on the full sample (including up to 10,074 cases and 103,164 controls) were only available for the 10,000 most robustly associated single nucleotide polymorphisms (SNPs). For the other SNPs, summary estimates were based on analyses excluding the 23andMe cohort (up to 4,890 cases and 20,405 controls). We therefore used summary statistics based on a combined version of the available data, with preference given to SNP statistics including the 23andMe cohort (denoted “Combined” dataset) and a window size spanning 2 Mb around the most robustly associated PCOS SNP based on P-value (19,24,25).

**Table 1.** Summary statistics for the top 14 single nucleotide polymorphisms associated with polycystic ovary syndrome from Day *et al.* (18). SNP: single nucleotide polymorphism; Chr: chromosome; Pos: position (hg19); EA: effect allele; NEA: non-effect allele; EAF: effect allele frequency; CI: confidence interval; P: P-value.

| SNP | Chr | Pos | EA | NEA | EAF | Estimate (95% CI) | P |
| --- | --- | --- | --- | --- | --- | --- | --- |
| rs2178575 | 2 | 213391766 | A | G | 0.15 | 1.18 (1.13-1.23) | $3.34 \times 10^{-14}$ |
| rs11031005 | 11 | 30226356 | C | T | 0.15 | 1.17 (1.12-1.23) | $8.66 \times 10^{-13}$ |
| rs804279 | 8 | 11623889 | A | T | 0.26 | 1.14 (1.10-1.18) | $3.76 \times 10^{-12}$ |
| rs11225154 | 11 | 102043240 | A | G | 0.09 | 1.20 (1.13-1.26) | $5.44 \times 10^{-11}$ |
| rs9696009 | 9 | 126619233 | A | G | 0.07 | 1.22 (1.15-1.30) | $7.96 \times 10^{-11}$ |
| rs13164856 | 5 | 131813204 | T | C | 0.73 | 1.13 (1.09-1.18) | $1.45 \times 10^{-10}$ |
| rs1784692 | 11 | 113949232 | T | C | 0.82 | 1.15 (1.10-1.21) | $1.88 \times 10^{-10}$ |
| rs7563201 | 2 | 43561780 | G | A | 0.55 | 1.11 (1.08-1.15) | $3.68 \times 10^{-10}$ |
| rs8043701 | 16 | 52375777 | T | A | 0.18 | 1.14 (1.09-1.18) | $9.61 \times 10^{-10}$ |
| rs1795379 | 12 | 75941042 | C | T | 0.76 | 1.12 (1.08-1.17) | $1.81 \times 10^{-09}$ |
| rs853854 | 20 | 31420757 | T | A | 0.50 | 1.10 (1.07-1.14) | $2.36 \times 10^{-09}$ |
| rs2271194 | 12 | 56477694 | A | T | 0.42 | 1.10 (1.07-1.14) | $4.57 \times 10^{-09}$ |
| rs10739076 | 9 | 5440589 | A | C | 0.31 | 1.12 (1.07-1.16) | $2.51 \times 10^{-08}$ |
| rs7864171 | 9 | 97723266 | G | A | 0.57 | 1.10 (1.06-1.13) | $2.95 \times 10^{-08}$ |

**Figure 1.**
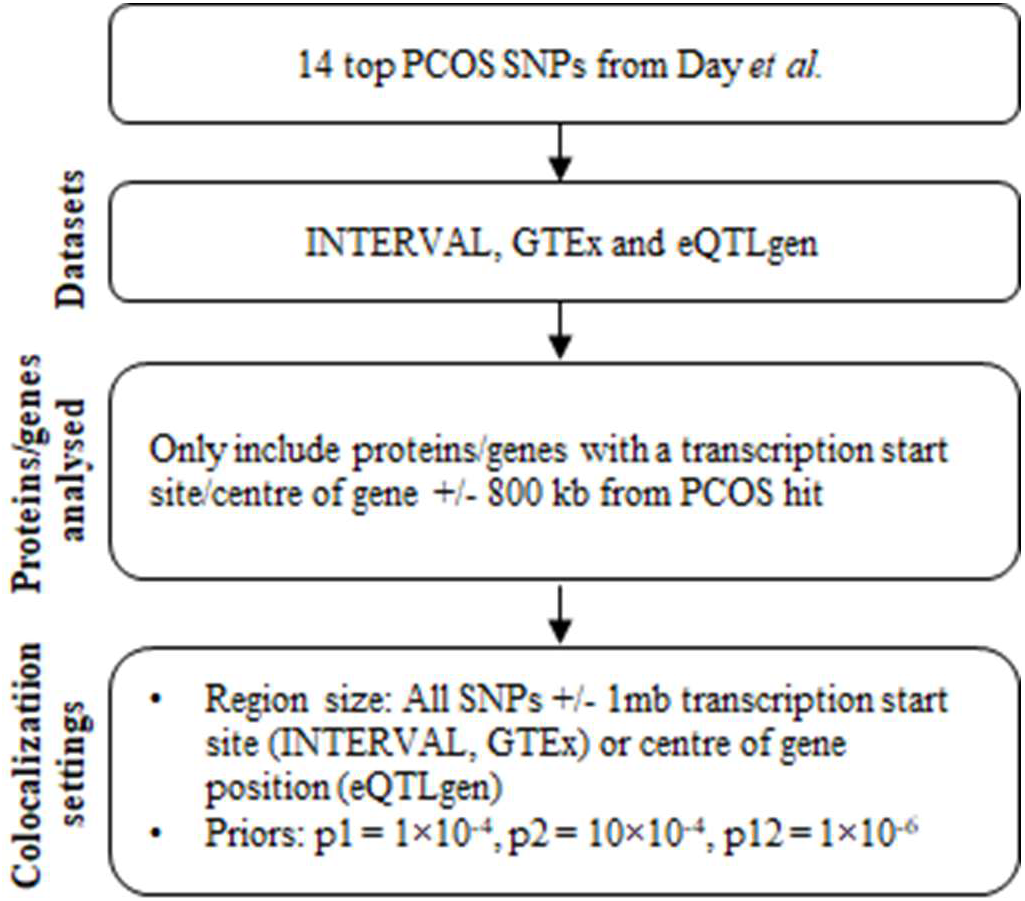
Study overview.

We identified seven proteins and genes with evidence of colocalization (posterior probability (PP) ≥ 0.75), including the protein FSH, and the genes *SUOX*, *ERBB3*, *IKZF4*, *RPS26*, *C8orf49*, and *ZFP36L2* (26,27). In addition, four genes (*RAD50*, *GDF11*, *NEIL2*, *C9orf3*) showed nominal evidence of colocalization (PP > 0.50) (Fig 2 and Supplementary Tables 1-3; for a detailed description of genes not discussed below see the Supplement and Supplementary Figures 1-9). Some of these genes and proteins, such as *RAD50*, had evidence of colocalization in only one tissue, whereas others, such as *RPS26* and *SUOX*, had evidence of colocalization in a large proportion of all tested tissues (Supplementary Tables 1-2). However, tissue sample size seemed to influence the evidence of colocalization and a large proportion of the colocalizing gene-tissue combinations used blood expression data from eQTLgen (sample size up to 31,684), but many of these analyses did not surpass the colocalization threshold using the smaller GTEx blood expression dataset (sample size up to 369) (Supplementary Table 2).

**Figure 2.**
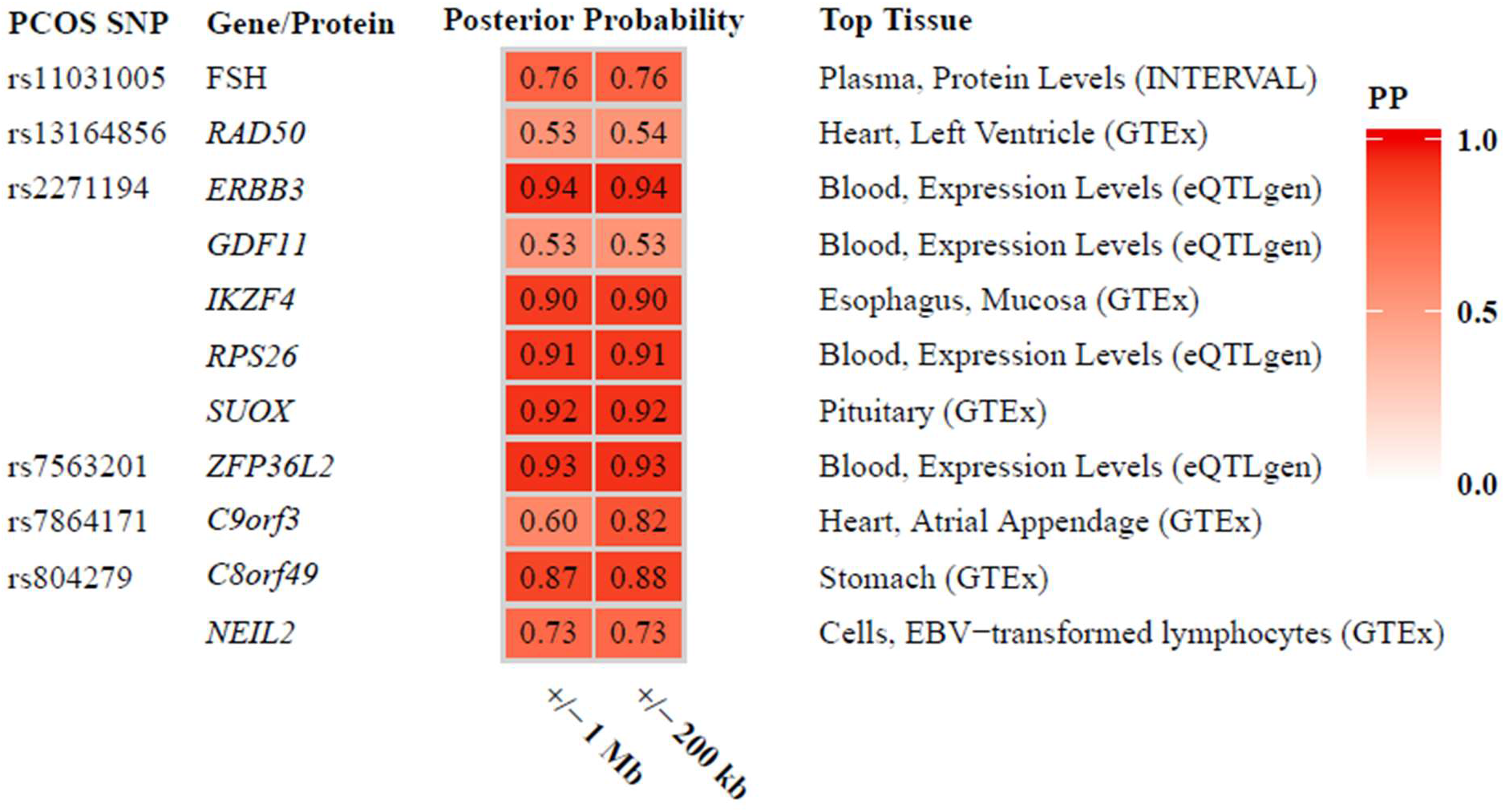
Posterior probabilities for genes and proteins with any evidence of colocalization. In the main approach, we used the Combined PCOS dataset and a region size spanning +/− 1 Mb. Only the results for the tissue with the highest posterior probability of colocalization in the main analysis are reported here (for full results and power calculations see Supplementary Table 1-3). Gene-tissue combinations with a posterior probability of colocalization >0.50 were seen as having some evidence in favour of colocalization, whereas the threshold for strong evidence was set at ≥0.75. PCOS, polycystic ovary syndrome; PP, posterior probability.

### Interaction-coloc analyses

Several genes and proteins had evidence of colocalization in some loci, which might be due to shared regulatory mechanisms (Fig 2). In addition, identification of true causal genes/proteins is dependent on tissue- and timepoint relevant QTL datasets, an inherent problem in colocalization analyses (19,28). We therefore suggest an exploratory approach, an “interaction-coloc”-analysis, to further query the evidence for each colocalizing gene/protein.

We reasoned that we could nuance the evidence of PCOS involvement for the colocalizing genes/proteins by assessing if other genes/proteins known to interact with them also had evidence of colocalization (Supplementary Figure 10). Specifically, if there is evidence of colocalization with PCOS for two genes/proteins known to interact with each other, this should in theory increase the likelihood of them and their affiliated pathway mediating the relationship with the disease (Fig 3). We therefore extracted protein-protein interaction data from Reactome (29) for the proteins and genes colocalizing (PP > 0.50) in our main analysis.

**Figure 3.**
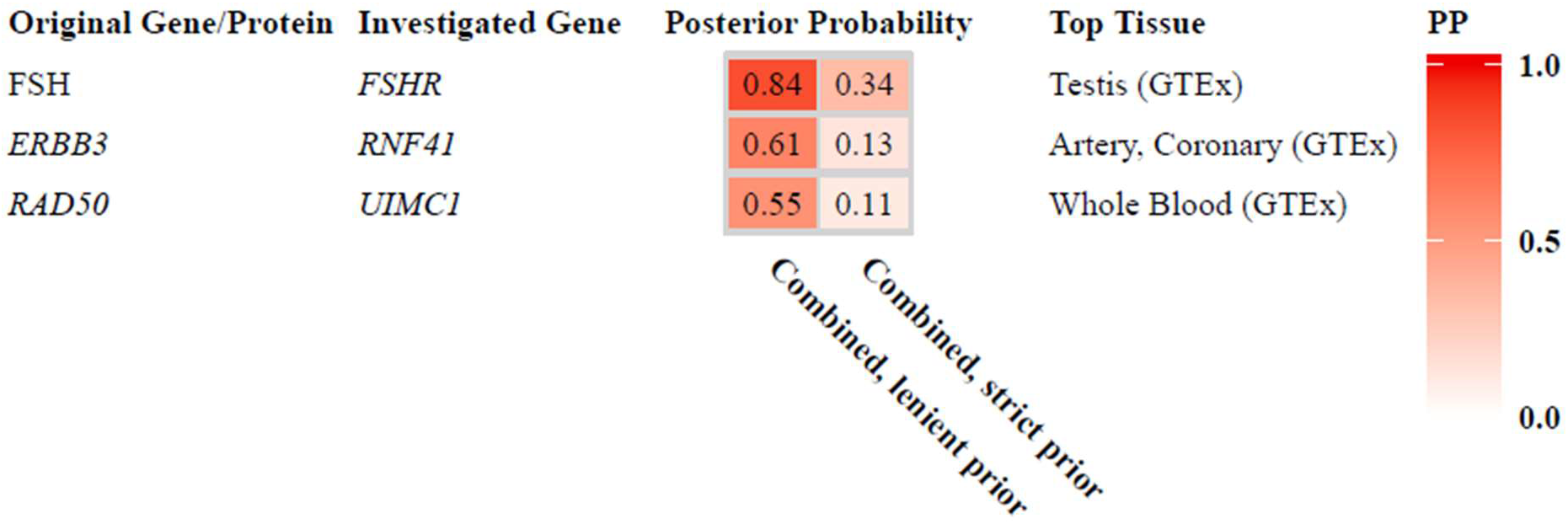
Posterior probabilities for genes with nominal evidence of colocalization in the interaction-coloc analyses. In the main approach, we used the Combined PCOS dataset and a set the prior probability of colocalization to p12 = 1×10^−5^. Sensitivity analyses included a more stringent prior probability of p12 = 1×10^−6^. Note that *RNF41* – implicated in the same pathway as *ERBB3* – was also located in the rs2271194 locus. PP, posterior probability.

We then performed colocalization for these “interactors” (including both their genes and any protein products) with PCOS risk. Using this approach, we found evidence of colocalization for *FSHR* expression (interacting with FSH), and nominal evidence of colocalization for *RNF41* (interacting with *ERBB3)* and *UIMC1* (interacting with *RAD50*) expression (Fig 3-5 and Supplementary Table 4) (29).

### Regulatory annotations and associations with other traits

Next, we analyzed phenome-wide associations (PheWAS) of the PCOS loci by characterizing their associations with other traits using public data (Supplementary Tables 5-10) (30). We also assessed regulatory evidence using Haploreg (31).

The colocalization results had highlighted circulating FSH as colocalizing at the rs11031005 locus (PP=0.76). We found that the rs11031005 C-allele was associated with both higher PCOS risk (OR 1.17, 95% CI 1.12-1.23, P=8.7×10^−13^) and lower FSH-levels (−0.166 standard deviations, standard error = 0.035, P = 2.0×10^−6^). In addition, rs11031005 was associated with several traits related to female hormonal regulation in the PheWAS look-up, with the two traits showing the most robust associations being length of menstrual cycle (P=1.2×10^−42^) and age at menopause (P=1.4×10^−15^) (Supplementary Table 5) (30,32).

Other PCOS loci seemed more pleiotropic – at the rs2271194 locus, the results supported colocalization for four genes (*ERBB3, IKZF4, RPS26*, and *SUOX*), as well as nominal evidence for *GDF11* (Fig 2, Supplementary Figures 1-5). The PheWAS of this locus highlighted associations with a range of different traits, including e.g. obesity, hematologic, and social traits (Supplementary Table 6) (30). Look-up of the PCOS SNP and its proxies (r2>0.8 in Europeans) in Haploreg (31) gave further evidence for a regulatory function acting in a many different cell-types, including the presence of enhancer and promoter marks, location in DNase hypersensitivity sites, and binding of e.g. RNA polymerase II and transcription factors (31,33–37).

### Sensitivity analyses and choice of priors

We performed several sensitivity analyses. Firstly, coloc uses SNP-associations to compute posterior probabilites (19), and association statistics are dependent on sample size. However, summary statistics for the entire PCOS sample (up to 10,074 cases and 103,164 controls) was only publicly available for the 10,000 most robustly associated SNPs. In contrast, full GWAS summary statistics were available for up to 4,890 cases and 20,405 controls (data based on analyses excluding the 23andMe cohort, denoted “Without-23” dataset). To ascertain similar sample sizes for all SNPs regardless of the strength of association, we therefore also performed colocalization using only the Without-23 PCOS dataset. Colocalization analyses using the Without-23 PCOS dataset generally had lower power (possible range 0-1, with a power >0.80 indicating strong power to determine colocalization) to detect colocalization, and generally a correspondingly lower PP of colocalization (Supplementary table 1-3) (24). For example, there was strong power and evidence for colocalization (power = 1.00 and PP = 0.93) between PCOS risk and expression of *ZFP36L2* at the rs7563201 locus using the Combined PCOS dataset, but considerably less power and PP using the Without-23 dataset (power = 0.28 and PP = 0.01).

Secondly, the number of SNPs included in the analysis can affect the PP of colocalization (25). We therefore also conducted analyses using a region size of +/− 200 kb for all three e/pQTL datasets (19,25), as well as approximately independent regions of linkage disequilibrium (38) in INTERVAL (performed in INTERVAL only since the other datasets did not provide genome-wide summary statistics) (39). In general, there was good consistency between all three window sizes (Figure 2, Supplementary tables 1-2).

Thirdly, we performed colocalization analysis using the software HyPrColoc to minimize the risk of software or coding errors (39). These results supported the main results (Supplementary Tables 1-2).

Finally, coloc requires specification of prior probabilities for both the likelihood that a SNP is associated with each trait (p1 and p2, respectively) and for the likelihood that a SNP is associated with both traits (p12). A previous study has shown that p1 = p2 = 1×10^−4^ is a reasonable setting in most scenarios, but the choice of p12 is more complex (25). We therefore decided to set p12 = 1×10^−6^ in the main analysis, corresponding to a stricter p12 than suggested (25) and stricter than the standard setting (19). For the interaction-coloc analyses, we used the standard coloc setting of p12 = 1×10^−5^, given a hypothesized greater likelihood of colocalization in these analyses, as well as p12 = 1×10^−6^ as a sensitivity analysis (Figure 3, Supplementary Table 4).

## Discussion

Our results highlight several genes and proteins that may have a role in PCOS development by using a Bayesian colocalization approach. We identify seven genes and proteins with strong and a further four genes and proteins with some evidence of colocalization, respectively. Several of these genes and proteins have links to the HPG axis and follicular development, further highlighting disruption of these processes as likely pathophysiological mechanisms in the disease. As the mediating genes for most of the genetic risk loci are still unclear (17,18), our results offer a potential to focus further functional follow-up studies on genes with a higher likelihood of being involved in PCOS pathophysiology.

Our results highlighted FSH (its beta-chain encoded by *FSHB*, located approximately 26 Kb from rs11031005 (33,40)) as a potential mediator at the rs11031005 locus. The results also implicated *ZFP36L2* at the rs7563201 locus. Female mice with a disruption in the *ZFP36L2* gene have disturbed oocyte maturation and ovulation, and its gene product has been implicated in regulation of LH-receptor levels (33,41). There is previous evidence for disruptions in gonadotropin signalling, specifically FSH and LH, being involved in PCOS pathophysiology (8,42). FSH and LH are crucial hormones for follicular development and ovulation (5,6,8), The two hormones share an alpha chain (encoded by *CGA* (33)), and disruption of *FSHB* has been associated with higher LH levels in both humans and mice (43,44). SNPs in the *FSHB* region have also been associated with levels of both LH and LH/FSH (45–47). It is thus possible that the PCOS association at the rs11031005 locus may partly be caused by altered *FSHB* expression affecting LH-levels, although the interaction-coloc evidence for involvement of the FSH-receptor also implies a direct role of FSH in the disease.

At the rs2271194 (at position 12:56477694 in GRCh 37 (48)) locus, two of the colocalizing genes – *ERBB3* and *RPS26* – are likely candidates for mediating PCOS risk based on the literature, with both of them connected to the HPG-axis (for a literature review of the other genes see Supplement). The gene *ERBB3* encodes a tyrosine-protein kinase receptor (Receptor tyrosine-protein kinase erbB-3) (33). *ERBB3* expression levels in granulosa cells vary over the estrous cycle in rats, with gonadotropins upregulating *ERBB3* expression and data suggesting an important role in follicular development (49,50). There was evidence of colocalization for *RNF41* (involved in regulation of Receptor tyrosine-protein kinase erbB-3 protein levels (33,51)) in the interaction-coloc analyses, but as the genes *ERBB3* and *RNF41* are in the same locus this cannot be regarded as additional evidence for *ERBB3*. The other likely candidate at the locus, *RPS26*, has been implicated in DNA damage response and female fertility (33,52,53). For example, oocyte-specific *Rps26*-knock-out mice have arrested oocyte growth, impaired follicle development, as well as poor response to gonadotropin stimulation (53), hence also implicating the HPG axis.

Another promising gene candidate is *RAD50*. The gene encodes DNA repair protein RAD50 (33), which together with MRE11 and another protein forms part of the MRE11 complex, which is involved in DNA damage response processes (54–59). Female mice with disruptions in the *Mre11* or *Rad50* genes have reduced fertility (55,59). It may be that the MRE complex affects oocyte elimination in the presence of DNA damage and thereby plays a part in follicular development and oocyte development (57). Even though our results only provided nominal evidence for involvement of *RAD50* in PCOS development, the evidence was strengthened by the interaction-coloc analyses that also gave nominal colocalization evidence for another gene (*UIMC1*) implicated in the same DNA repair processes as the MRE11 complex (33,60).

Importantly, shared regulatory mechanisms between e.g. different genes and tissues can result in several gene/protein and tissue combinations colocalizing. However, it is unlikely that all of them are involved in disease development – indeed, the true mediating gene and tissue combination may not even have been investigated in the analyses. Therefore, while colocalization can highlight genes and proteins that are more likely to be involved in PCOS pathophysiology, results should be seen as hypothesis-generating rather than definitive evidence of a causal role.

Whereas some genes exhibit more tissue-specific effects, others have similar effects in a range of tissues (20,61,62). We assessed colocalization using datasets including a wide range of tissue types (e.g. GTEx (20)) and datasets with large sample sizes (e.g. eQTLgen (21)), which should increase the chance of identifying colocalizing genes and proteins.

We also investigated if genes/proteins that may interact with the originally identified genes/proteins provided additional evidence of their involvement in PCOS pathophysiology. This is a novel approach, but whereas it in theory should provide a more independent confirmation of a gene/protein being involved in the disease, the results should be interpreted with caution. Some of the originally identified genes and proteins had many known interactors and others none, resulting in differing possibilities to identify colocalization. In addition, even though the interaction-coloc analysis delivered plausible results and presents a possible extension of colocalization methodology, it has not been validated.

There are also caveats with our study. Firstly, if the causal SNP (or a proxy) is altering the coding sequence of a tested protein, it may cause false positive results through changed aptamer binding. Secondly, ancestral heterogeneity could potentially bias results due to different LD-structure (19), even though all datasets primarily consisted of participants of European decent (20–22,63). Thirdly, the protein and expression datasets included both men and women (20–22,63), whereas the PCOS GWAS (18) was performed in women only. If associations between genotypes and expression/protein levels differ between the sexes, it could bias results. Finally, *coloc* assumes a single causal variant per locus (19). Accordingly, loci with multiple SNPs independently associated with either the disease or the intermediate trait risk may result in false negative results (19).

## Conclusion

In summary, our results highlight potential mediating genes and proteins for almost a third of PCOS risk loci. Several of these genes and proteins have links to the HPG axis and follicular development, including the hormone FSH and the genes *ZFP36L2, ERBB3, RPS26*, and *RAD50*. In combination with previous studies that have indicated these genes as being involved in physiologic processes associated with PCOS, these genes may be of particular interest for further functional follow-up.

## Materials and Methods

### Data on Polycystic ovary syndrome

We obtained GWAS summary statistics for PCOS from Day *et al.* (18). In the study, 14 genome-wide significant loci were identified in up to 10,074 cases and 103,164 controls of European ancestry. Public summary statistics were available for the full sample for the 10,000 most robustly associated SNPs, and for all SNPs from analyses excluding the 23andMe cohort (resulting in a sample size of up to 4,890 cases and 20,405 controls). To maximize power, we used a combined version of these two datasets as our main dataset (denoted “Combined” dataset), with preference given to data from the top 10,000 SNPs dataset. As a sensitivity analysis, we also performed all analyses using the all-SNP dataset where the 23andMe cohort had been excluded (denoted “Without-23” dataset), to have roughly the same sample size for all SNPs. We then excluded SNPs found to be duplicated by position, missing relevant data, or indels. Genetic variants were matched to rsIDs using the file “All_20180423.vcf.gz”, available at ftp://ftp.ncbi.nih.gov/snp/organisms/human_9606_b151_GRCh37p13/VCF/ (48).

### Quantitative trait loci datasets

We used publicly available protein and expression genetic association data from the INTERVAL study (22,23), the GTEx consortium (20), and the eQTLgen consortium (21).

pQTL data were taken from the INTERVAL study, which had performed GWASs for 2,994 unique plasma proteins (3,283 measured aptamers) in 3,301 blood donors of European ancestry (22). For GTEx, we used data from version 7, which contains cis-eQTL data for between 80-491 samples in 48 different tissues (20,63). Expression had been measured post-mortem, with ~85% of donors being of European (“White”) ancestry in the whole sample (63). Lastly, the eQTLgen Consortium had performed cis- and trans-eQTL analysis in up to 31,684 individuals, predominantly of European ancestry (21). Both cis-associations, containing SNPs within 1 Mb from the center of the gene, and trans-assocations, containing SNPs over 5 Mb from the center of the gene, are publicly available (21). For all these datasets, we then excluded SNPs that were duplicated by position, missing relevant data, or indels.

### Colocalization analyses

#### Coloc

We applied coloc (19), a Bayesian test for colocalization to evaluate the probability of a shared causal signal between each PCOS hit and each p/eQTL. We performed colocalization using the coloc.abf() function in the coloc R package, applying it to cis-genes using up to three different region sizes depending on QTL dataset. Gene positions and transcription start sites were determined using GRCh 37 and the biomaRt R package where needed (64,65).

For GTEx and eQTLgen, cis-association statistics were only available for SNPs within 1 Mb of the transcription start site and the centre of the gene, respectively (20,21). We therefore only included genes and proteins with a transcription start site or centre of gene +/− 800 kb of the top PCOS SNP (by P-value) for all three QTL datasets, to ascertain that we had a sufficiently large region on both sides of the association peak to determine colocalization. We further analysed two different region sizes in GTEx and eQTLgen – the entire 2 Mb cis-region available in these datasets in the main analysis and +/− 200 kb of the top SNP as a sensitivity analysis. For GTEx, we only performed the analysis if the top SNP had been analyzed for computational reasons. For INTERVAL (22), we evaluated three different region sizes – +/− 1 Mb and +/− 200 kb of the top SNP, as well as the top SNP’s “independent region” (19,24,39,66). Independent regions were defined as the approximately independent regions of linkage disequilibrium in Europeans, as computed by Berisa *et al.* (38).

We set the prior probabilities to p1 = 1×10^−4^, p2 = 10×10^−4^, and p12 = 1×10^−6^ (more stringent than default) (19,25). Minor allele frequencies from the PCOS dataset were used in all coloc analyses. For the number of cases and total sample size, we supplied 10,074 and 113,238 for the Combined dataset (albeit this would be smaller for the SNPs that were not in the top 10,000 SNPs dataset) and 4,890 and 25,295 for the sensitivity-analysis using the Without-23 PCOS dataset (the dataset with estimates based on approximately equal sample sizes for each SNP). For INTERVAL and GTEx, we used the sample size reported for each tissue and dataset. For eQTLgen we supplied the average sample size for the included SNPs. As the eQTLgen summary statistics did not include effect estimates and standard errors, we let the coloc.abf() function approximate effect estimates from the P-values for this dataset (19).

Briefly, coloc evaluates the PP for five different hypotheses, which in this study correspond to:

- H_0_: No causal association with either PCOS or the protein/gene
- H_1_: Causal association with PCOS but not the protein/gene
- H_2_: Causal association with the protein/gene but not PCOS
- H_3_: Causal associations with both PCOS and the protein/gene, but with two separate causal SNPs
- H_4_: Causal association with both PCOS and the protein/gene, with a shared causal SNP (19)

Studies use different thresholds to evaluate whether there is evidence of a shared causal variant (H_4_), but the PP of colocalization can be seen as a numerical value of the certainty of the result (19,26,66–68). Since we performed colocalization as a hypothesis-generating approach, all analyses with a PP >0.50 were seen as having nominal evidence of colocalization and analyzed further. A PP just above >0.50 should be regarded with caution (19), and we set the threshold for strong evidence of colocalization at PP ≥0.75 (26,27). We also computed the power for detecting colocalization for the results with any evidence of colocalization as the sum of the PPs for hypothesis 3 (no colocalization) and hypothesis 4 (colocalization) (24).

#### HyPrColoc

To ascertain robustness, we also computed the posterior probability of colocalization using HyPrColoc (39), a recently developed extension of coloc (19). We used a similar approach as for coloc, but only using the larger region sizes of 1 Mb for all three QTL datasets, as well as the independent regions for INTERVAL. Default priors (prior.1 = 1×10^−4^ and prior.2 = 0.98) were used, whereas we set both the regional and alignment probability thresholds to 0.8 (more stringent than default) (39). As eQTLgen only provided Z-scores, we estimated betas and SEs using the formulas:

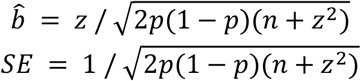

Where *z* is the Z-score, *p* is the minor allele frequency in the eQTLgen dataset and *n* is the sample size (21,69).

#### Protein-protein interaction follow-up analyses using coloc

To identify genes/proteins that interact with the primarily identified genes/proteins, we downloaded data with protein-protein interactions in humans (available at https://reactome.org/download/current/interactors/reactome.homo_sapiens.interactions.tab-delimited.txt) from Reactome (29). Genes listed as part of proteins interacting with any of our associated genes, and with ensembl gene identifiers, were extracted. For FSH, we only extracted interactions listed for the beta subunit (encoded by *FSHB*), since the alpha subunit (encoded by *CGA*) forms part of other hormones as well (70). We then extracted information of uniprot-identifiers, ensembl gene identifiers, gene positions and transcription start sites using GRCh 37 and the biomaRt R package to map between different datasets (64,65). We only included transcripts listed with a numeric autosomal chromosome and with information available in biomaRt. We included SNPs within +/− 1 Mb from the average transcription start site in the colocalization analyses using INTERVAL dataset (22), for the other datasets all available SNPs were used. We then applied coloc (19), using +/− 1 Mb region sizes. As the genes and proteins in the interaction-coloc analyses already had evidence of protein-protein interactions with the genes identified in the main analyses, we considered the prior probability of colocalization higher and thus used a more lenient prior probability of colocalization than in the main analysis (p12 = 1×10^−5^, which is the same as the default setting in coloc (19)).

### PheWAS and in-silico investigations

We followed up colocalizing regions with assessing PheWAS data for the top PCOS SNP using the Open Target Genetics platform (30). The significance threshold for a PheWAS association on the Open Targets Genetics platform is approximately P<1×10^−5^ (based on visual inspection of the plotted threshold, which corresponds to a Bonferroni-correction of the number of investigated traits (30)). We further corrected for the six SNPs we investigated and set the threshold to P<1.7×10^−6^ (1×10^−5^ corrected for six SNPs). We also investigated the evidence for regulatory mechanisms for the colocalizing PCOS regions and the top SNP using Haploreg v4.1 (31).

### Software

Analyses and plots were done using R versions 3.5.1 and 3.4.3 (71), bash version 4.1.2(2) (72), awk (73), and R packages coloc (19), hyprcoloc (39), LocusCompareR (74), tidyr (75), data.table (76), plyr (77), devtools (78), and ggplot2 (79).

**Figure 4A and 4B.**
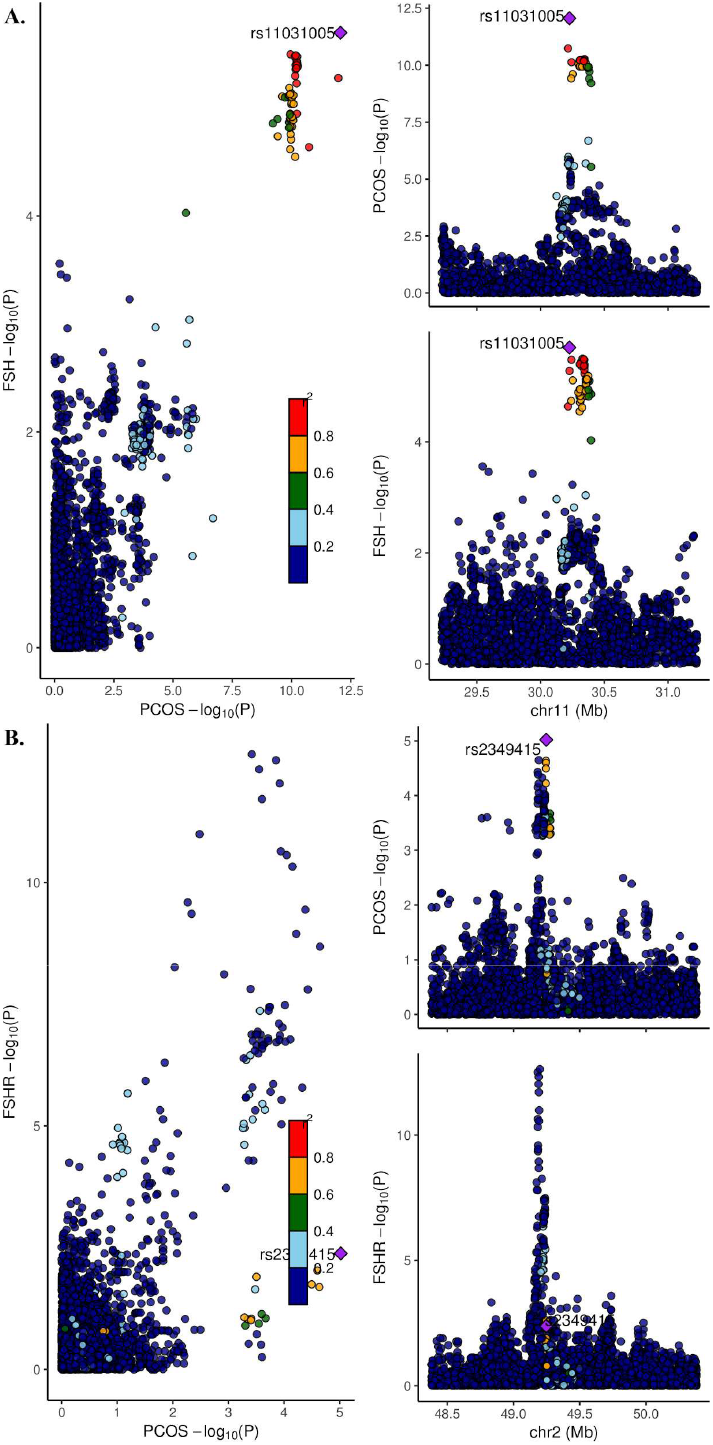
Associations between genetic variants and PCOS risk, using Combined PCOS dataset, +/− 1 Mb region sizes for (A) FSH protein levels in blood (B) *FSHR* expression levels in testis. In each plot, each dot is a genetic variant. The SNP with the most significant P-value for PCOS is marked, with the other SNPs colour-coded according to linkage disequilibrium (r^2^) in Europeans with the lead variant. SNPs with missing linkage disequilibrium information are also coded dark blue. In the left panels, -log10 P-values for associations with PCOS risk are on the x-axes, and -log10 P-values for associations with the protein/transcript levels on the y-axes. On the right panels, genomic positions are on the x-axes, and the y-axes show -log10 P-values for PCOS on the upper panel and -log10 P-values with the protein/expression levels on the lower panel for the corresponding region. FSH, follicle stimulating hormone; PCOS, polycystic ovary syndrome; SNP, single nucleotide polymorphism.

**Figure 5A and 5B.**
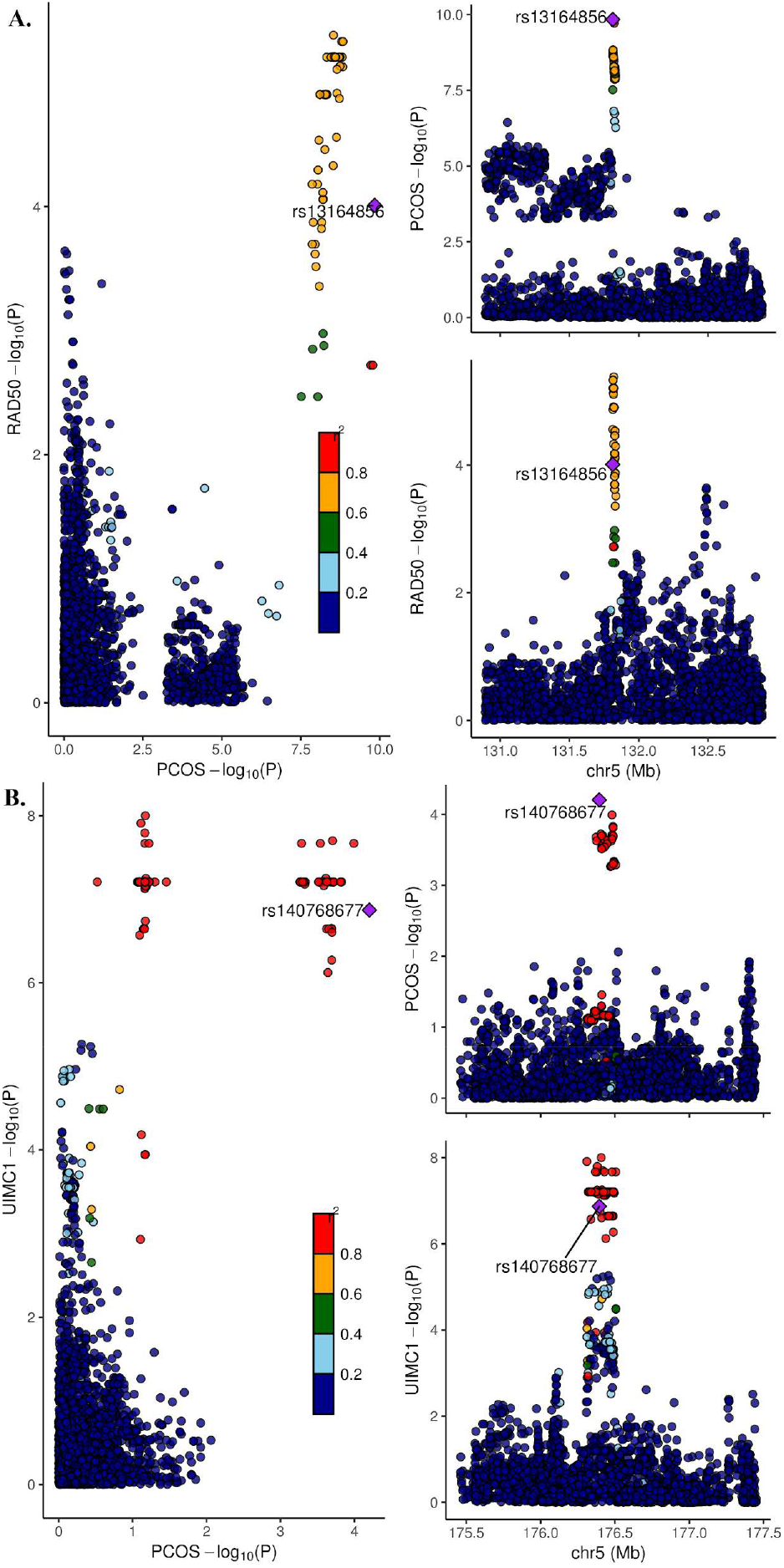
Associations between genetic variants and PCOS risk, using Combined PCOS dataset, +/− 1 Mb region sizes for (A) *RAD50* expression levels in left ventricle of the heart (B) *UIMC1* expression levels in blood (GTEx) In each plot, each dot is a genetic variant. The SNP with the most significant P-value for PCOS is marked, with the other SNPs colour-coded according to linkage disequilibrium (r^2^) in Europeans with the lead variant. SNPs with missing linkage disequilibrium information are also coded dark blue. In the left panels, -log10 P-values for associations with PCOS risk are on the x-axes, and -log10 P-values for associations with the expression levels on the y-axes. On the right panels, genomic positions are on the x-axes, and the y-axes show -log10 P-values for PCOS on the upper panel and -log10 P-values with the expression levels on the lower panel for the corresponding region. PCOS, polycystic ovary syndrome; SNP, single nucleotide polymorphism.

## Data availability

The PCOS GWAS summary statistics are available at https://www.repository.cam.ac.uk/handle/1810/283491 (18). The GTEx version 7 data are available at https://gtexportal.org/ (20). Effect allele frequencies for GTEx were taken from the files “GTEx_V7_cis_eqtl_summary.tar.gz (hg19)” (downloadable at http://cnsgenomics.com/software/smr/#DataResource). Independent regions as per Berisa *et al.* (38) can be accessed at https://bitbucket.org/nygcresearch/ldetect-data/downloads/. The summary statistics from the INTERVAL study is available at https://www.phpc.cam.ac.uk/ceu/proteins/ (22). Data from the eQTLgen consortia can be accessed at https://molgenis26.gcc.rug.nl/downloads/eqtlgen/cis-eqtl (21). Human protein-protein interactions from Reactome pathways is available at https://reactome.org/download/current/interactors/reactome.homo_sapiens.interactions.tab-delimited.txt. The PheWAS data were downloaded from the Open Targets Genetics website https://genetics.opentargets.org (30). In-silico functional investigations were done using Haploreg v4.1 at https://pubs.broadinstitute.org/mammals/haploreg/haploreg.php (31). Individual-level data from UK Biobank cannot be shared publicly because of confidentiality but is available from the UK Biobank (https://www.ukbiobank.ac.uk/) for researchers who meet the criteria for access to confidential data. The UK Biobank has a Research Tissue Bank approval (Research Ethics Committee reference 16/NW/0274, this study’s application ID 11867).

## Acknowledgements

We thank the PCOS Consortium. The Genotype-Tissue Expression (GTEx) Project was supported by the Common Fund of the Office of the Director of the National Institutes of Health, and by NCI, NHGRI, NHLBI, NIDA, NIMH, and NINDS. Computation used the Oxford Biomedical Research Computing (BMRC) facility, a joint development between the Wellcome Centre for Human Genetics and the Big Data Institute supported by Health Data Research UK and the NIHR Oxford Biomedical Research Centre. The views expressed are those of the author(s) and not necessarily those of the NHS, the NIHR or the Department of Health. We thank the UK Biobank (http://www.ukbiobank.ac.uk/; application id 11867).

## Abbreviations

eQTL: expression quantitative trait locus
FSH: follicle-stimulating hormone
GnRH: gonadotropin-releasing hormone
GTEx: Genotype-Tissue Expression project
GWAS: genome-wide association study
HPG: hypothalamic-pituitary-gonadal
LH: luteinizing hormone
PCOS: polycystic ovary syndrome
PheWAS: phenome-wide association study
PP: posterior probability
pQTL: protein quantitative trait locus
SNP: single nucleotide polymorphism

