## Supplementary Material for "Colocalization highlights genes in hypothalamic–pituitary–gonadal axis as potentially mediating polycystic ovary syndrome risk"

#### Table of Contents


Evaluation of performing colocalization using the top 10,000 SNPs PCOS dataset 3

Linkage-disequilibrium reference panel for the conditional analyses 3

Conditional analysis of *FSHR* expression 3

Genes with evidence of colocalization only in sensitivity analyses 4

Literature review of genes 4

- rs2271194 and *IKZF4* 4
- rs2271194 and *GDF11* 4
- rs2271194 and *SUOX* 4
- rs7864171 and *C9orf3* 5
- rs804279 and *C8orf49* 5
- rs804279 and *NEIL2* 5
- rs10739076 and *RCL1* 5
- rs11031005 and *ARL14EP* 5
- rs11225154 and *TMEM123* 6
- rs2271194 and *ZC3H10* 6

Supplementary Figures 7

- Supplementary Figure 1A and 1B. Associations between genetic variants and PCOS risk, using Combined PCOS dataset, +/- 1 Mb region sizes for (A) *ERBB3* expression levels in blood (eQTLgen) (B) *RNF41* expression levels in coronary artery (GTEx) 7
- Supplementary Figure 2. Associations between genetic variants and PCOS risk, using Combined PCOS dataset, +/- 1 Mb region sizes for *GDF11* expression levels in blood (eQTLgen) 8
- Supplementary Figure 3. Associations between genetic variants and PCOS risk, using Combined PCOS dataset, +/- 1 Mb region sizes for *IKZF4* expression levels in esophagus (mucosa) 9
- Supplementary Figure 4. Associations between genetic variants and PCOS risk, using Combined PCOS dataset, +/- 1 Mb region sizes for *RPS26* expression levels in blood (eQTLgen) 10
- Supplementary Figure 5. Associations between genetic variants and PCOS risk, using Combined PCOS dataset, +/- 1 Mb region sizes for *SUOX* expression levels in pituitary 11
- Supplementary Figure 6. Associations between genetic variants and PCOS risk, using Combined PCOS dataset, +/- 1 Mb region sizes for *ZFP36L2* expression levels in blood (eQTLgen) 12
- Supplementary Figure 7. Associations between genetic variants and PCOS risk, using Combined PCOS dataset, +/- 1 Mb region sizes for *C9orf3* expression levels in atrial appendage of the heart 13
- Supplementary Figure 8. Associations between genetic variants and PCOS risk, using Combined PCOS dataset, +/- 1 Mb region sizes for *C8orf49* expression levels in stomach 14

Supplementary Figure 9. Associations between genetic variants and PCOS risk, using Combined PCOS dataset, +/- 1 Mb region sizes for *NEIL2* expression levels in EBV-transformed lymphocytes 15

Supplementary Figure 10. Rationale for investigating the evidence for colocalization between PCOS and the protein/expression levels of other genes interacting with the originally identified genes. 16

Supplementary Figure 11A and 11B. Associations between genetic variants and PCOS risk, using Combined PCOS dataset for *CAPS2* expression levels in transverse colon using (A) Top 10,000 SNPs PCOS dataset (B) Combined PCOS dataset 17

Supplementary Figure 12A and 12B. Associations between genetic variants and PCOS risk, using Combined PCOS dataset, +/- 1 Mb region sizes for (A) *FSHR* expression levels in testis using estimates conditioned on rs2349415 (B) *FSHR* expression levels in testis using estimates conditioned on rs4953650 18

Supplementary Figure 13A and 13B. Associations between genetic variants and PCOS risk, using Combined PCOS dataset for *RCL1* expression levels in blood (eQTLgen) using (A) +/- 200 kb region size (B) +/- 1 Mb region size 20

References 22

#### Evaluation of performing colocalization using the top 10,000 SNPs PCOS dataset

We first performed colocalization using only the 10,000 most robustly associated single nucleotide polymorphisms (SNPs) (based on analyses including the 23andMe cohort) dataset, in addition to using the Combined polycystic ovary syndrome (PCOS) dataset and the Without-23 dataset. We used the same settings as in the main colocalization analysis (see main article), and the results using the top 10,000 SNPs datasets were largely comparable to those using the Combined PCOS dataset. However, we realized that this approach could potentially yield false positive results.

One example of such a potential false positive is the colocalization analysis of *CAPS2* expression levels in transverse colon and PCOS risk (Supplementary Figure 11 and Supplementary Table 2). In the colocalization analysis using +/- 1 Mb region sizes, there was nominal evidence in favour of colocalization using the top 10,000 SNPs PCOS dataset (posterior probability (PP)=0.56), but no evidence using the Combined or Without-23 dataset (PP=0 for both datasets).

Visual inspection of the region indicated that the light-blue SNPs ($r^2$ between 0.2 and 0.4 with the top PCOS SNP) associated with *CAPS2* expression levels around -log10 P-values of 7.5-10 in Supplementary Figure 11B likely offer evidence against colocalization in the analysis using the Combined PCOS dataset. However, as these SNPs are not strongly associated with PCOS risk, they are excluded in the analysis using only the top 10,000 SNPs dataset (Supplementary Figure 11A).

We realized that if the two traits have two distinct causal variants in linkage disequilibrium (LD) with each other, pre-selecting SNPs based on association with PCOS risk could result in the top expression quantitative trait loci (QTLs) SNPs being excluded from the analysis. Thus, SNPs providing evidence against colocalization would not be included, which could yield false positive results. We have therefore not presented the colocalization results for the top 10,000 SNPs in the main section of the paper, but have included them in Supplementary Table 2 for full transparency.

#### Linkage-disequilibrium reference panel for the conditional analyses

We constructed an LD-reference panel using a subset of the UK Biobank (1) and the imputed data for use in the analyses for computing conditional estimates. UK Biobank has a Research Tissue Bank approval (Research Ethics Committee reference 16/NW/0274, this study's application ID 11867), and all participants gave informed consent. Briefly, we extracted a random subsample of 29,454 female participants in the "White British ancestry" subset, after excluding individuals that had withdrawn consent, mismatch between self-reported and genetically inferred sex, sex-chromosome aneuploidy, reported incompatible ancestries in different assessments, that were related to other individuals in the UK Biobank to a third degree or higher, heterozygosity or missingness outliers, or not included in the autosome phasing or in the kinship calculations (2). Genotype dosages were converted to best-guess genotypes using a hard-call threshold of 0.1. We excluded SNPs with imputation info score $\leq 0.3$, minor allele frequency <0.01%, Hardy-Weinberg equilibrium exact test $P<1\times10^{-6}$, or genotype missing call rate >0.05. Analyses were done in plink versions 1.90b3 and 2.00a-20170724 (3).

#### Conditional analysis of *FSHR* expression

As the visual inspection of the *FSHR*-region (Figure 4 in the main article) indicated that there may be two or more independent eQTLs for *FSHR* expression in testis in the region, we performed colocalization analyses using estimates conditioned on top variants to determine which peak might be driving the colocalization. We first performed clumping on the *FSHR*-locus for PCOS (options --

clump-p1 0.0001, --clump-r2 0.05, and with --clump-kb spanning the entire locus), using the UK Biobank LD-reference panel. This identified two independent SNPs (rs2349415 and rs4953650, $r^2$=0.04 in Europeans using 1000 Genomes (4,5)). We then computed estimates conditioned on each of these top SNPs using GCTA version 1.91.4 (option --cojo-cond) for both the *FSHR* eQTL dataset in testis and the Combined PCOS dataset (6,7). Information on effect allele frequencies for GTEx were taken from the file "GTEx_V7_cis_eqtl_summary.tar.gz (hg19)" (downloadable at http://cnsgenomics.com/software/smr/#DataResource).

The colocalization analyses indicated that the colocalization was driven by the rs4953650 peak, as using estimates conditioned on rs2349415 yielded a colocalization PP of 0.78, whereas there was no evidence of colocalization when conditioning on rs4953650 (PP=0.02) (Supplementary Table 4, Supplementary Figure 12).

#### Genes with evidence of colocalization only in sensitivity analyses

Four genes (*RCL1*, *TMEM123, ARL14EP*, and *ZC3H10*) had evidence of colocalization only when using other coloc settings than the main approach, i.e. in the sensitivity analyses only (Supplementary Tables 1-2). We investigated these loci further, but the evidence of colocalization provided by visual inspection was in general weak and the genes had only weak support of a role in PCOS pathophysiology in the literature. For the interested reader, we have included a brief literature review of the genes below, together with those genes without detailed description in the main section of the paper.

#### Literature review of genes

##### rs2271194 and *IKZF4*

*IKZF4* encodes the protein Zinc finger protein Eos (8), which plays a role in gene regulation in T-regulatory cells and the immune response (9). Deletions of Eos in T-regulatory cells in mice induces autoimmunity (10). Whereas the posterior probability for colocalization was high, there is little evidence in the literature to support a role for the gene in PCOS pathophysiology at present (Supplementary Tables 1-3, Supplementary Figure 3).

##### rs2271194 and *GDF11*

The protein Growth/differentiation factor 11 (GDF11) has been implicated in several mammalian developmental processes (11–13), including adipogenesis (14) and pancreatic β-cell development (15). The GDF11 protein has also been shown to improve β-cell function in cells and islets from mice (16). Both follicle stimulating hormone (FSH) and the GDF11 protein are regulated by follistatin (17–21), and there is evidence for follistatin levels being higher in PCOS patients (22). In addition, mice treated with the androgen dehydroepiandrosterone have down-regulated expression levels of *GDF11* in the ovary (23). It is possible that increased risk at the rs2271194 locus is mediated through the effect of the GDF11 protein on adipogenesis and β-cell function and subsequent effects on e.g. insulin metabolism, or through yet unknown pathways in the ovary regulated by follistatin and/or androgen levels. However, more studies are needed to determine if GDF11 is involved in PCOS pathophysiology and, if so, the mechanistic pathways.

##### rs2271194 and *SUOX*

*SUOX* encodes mitochondrial sulfite oxidase (8), and it may be differentially expressed in oocytes from old versus young mice (24,25). The colocalization evidence for *SUOX* was comparable to the other genes in the rs2271194 PCOS risk locus (Supplementary tables 1-2). Whereas visual inspection of the associations between PCOS risk and *SUOX* expression levels supported colocalization, is is unclear if and how *SUOX* might affect PCOS risk (Supplementary Figure 5).

**rs7864171 and *C9orf3***

*C9orf3*, or *AOPEP* as the gene is also referred to, encodes Aminopeptidase O (8). Although the gene region has been linked to atrial fibrillation (26) and DNA methylation changes in twins born after in-vitro fertilization (27), the function of *C9orf3* is unclear. Our analyses indicated only moderate evidence of colocalization between *C9orf3* gene expression and PCOS risk (Supplementary Table 1-3 and Supplementary Figure 7).

**rs804279 and *C8orf49***

*C8orf49* is a long non-coding RNA (8,28). Very little is known about the gene, although its expression levels were included in a recent expression signature for prognosis of endometrial cancer (28). The evidence for colocalization was relatively high using the Combined PCOS dataset regardless of region size, but only for expression levels in stomach. The evidence of colocalization between *C8orf49* expression levels and PCOS risk in stomach diminished upon use of the Without-23 dataset, in spite of high power, wherefore the results should be interpreted with caution (Supplementary tables 1-3, Supplementary figure 8).

**rs804279 and *NEIL2***

The gene *NEIL2* codes for the protein Endonuclease 8-like 2 (8). The protein is a base excision repair gene important for long-term genomic maintenance (29) and its expression is prognostic for resistance to endocrine therapy in estrogen receptor positive breast cancer (30). However, *Neil2* knock-out mice have normal fertility (29,31). Whereas DNA damage repair is crucial for normal oocyte development (32), one study indicated undetectable levels of *NEIL2* in human oocytes and blastocysts (33). However, *NEIL2* has also been implicated in DNA demethylation together with tet3 (34), and female mice with tet3-depletion have reduced fecundity and their oocytes a reduced ability to reprogram somatic cells (35). In summary, more studies are needed to determine if *NEIL2* has a role in PCOS pathophysiology (Supplementary tables 1-3, Supplementary Figure 9).

**rs10739076 and *RCL1***

*RCL1* encodes the protein RNA 3'-terminal phosphate cyclase-like protein (8). The protein has been implicated in RNA processing (36), however not much else is known. In our colocalization analyses, there was only evidence for colocalization for the Combined PCOS dataset and blood expression levels in the +/- 200 kb region (Supplementary Table 1). Comparing the plots using the +/- 1 Mb and +/- 200 kb region sizes illustrates that part of the reason likely is a strong, second association peak for *RCL1* expression not present in the +/- 200 kb region (Supplementary Figure 13). Still, visual inspection of the smaller region size reveals that there only seems to be a single shared SNP moderately associated with *RCL1* expression. Taken together, there is only weak evidence for colocalization between PCOS risk and *RCL1* gene expression.

**rs11031005 and *ARL14EP***

The gene *ARL14EP* encodes the protein ARL14 effector protein (8). There is some evidence for the ARL14 effector protein being involved in MHC-class II molecule transportation (37), but it is unclear if it has any other function. There was evidence for *ARL14EP* colocalizing with PCOS risk, however primarily in testis and when using the Without-23 dataset. The PP was less than 0.50 when using the Combined PCOS dataset regardless of region size. Given the available data, the evidence points to FSH rather than *ARL14EP* mediating the PCOS risk at the rs11031005 PCOS risk locus.

**rs11225154 and *TMEM123***

Porimin, also known as Transmembrane protein 123 and encoded by *TMEM123*, is a transmembrane protein that has been implicated in cell death (8). Whereas *TMEM123* seems to be expressed in cumulus cells in the oocytes of PCOS patients (http://ivf.gxbsidra.org/dm3/geneBrowser/show/4000030, (38)), there is otherwise little evidence to link it to PCOS at present. The colocalization evidence for *TMEM123* was weak, with the highest PP in any coloc analysis 0.54 in brain cortex (Supplementary tables 1-2). In summary, there is little evidence to support a role of *TMEM123* in PCOS pathophysiology given our current knowledge.

**rs2271194 and *ZC3H10***

Few studies have investigated the function of *ZC3H10* (encoding the protein Zinc finger CCCH domain-containing protein 10 (8)), but it has been suggested that the gene is an important regulator of mitochondrial energy metabolism (39). In addition, a loss-of-function mutation in humans has been associated with metabolic phenotypes, including higher body mass index and fasting glucose (39). In contrast to the other genes in the rs2271194 PCOS risk locus, the evidence for colocalization between *ZC3H10* expression and PCOS risk was weak (Supplementary tables 1-2).

### Supplementary Figures

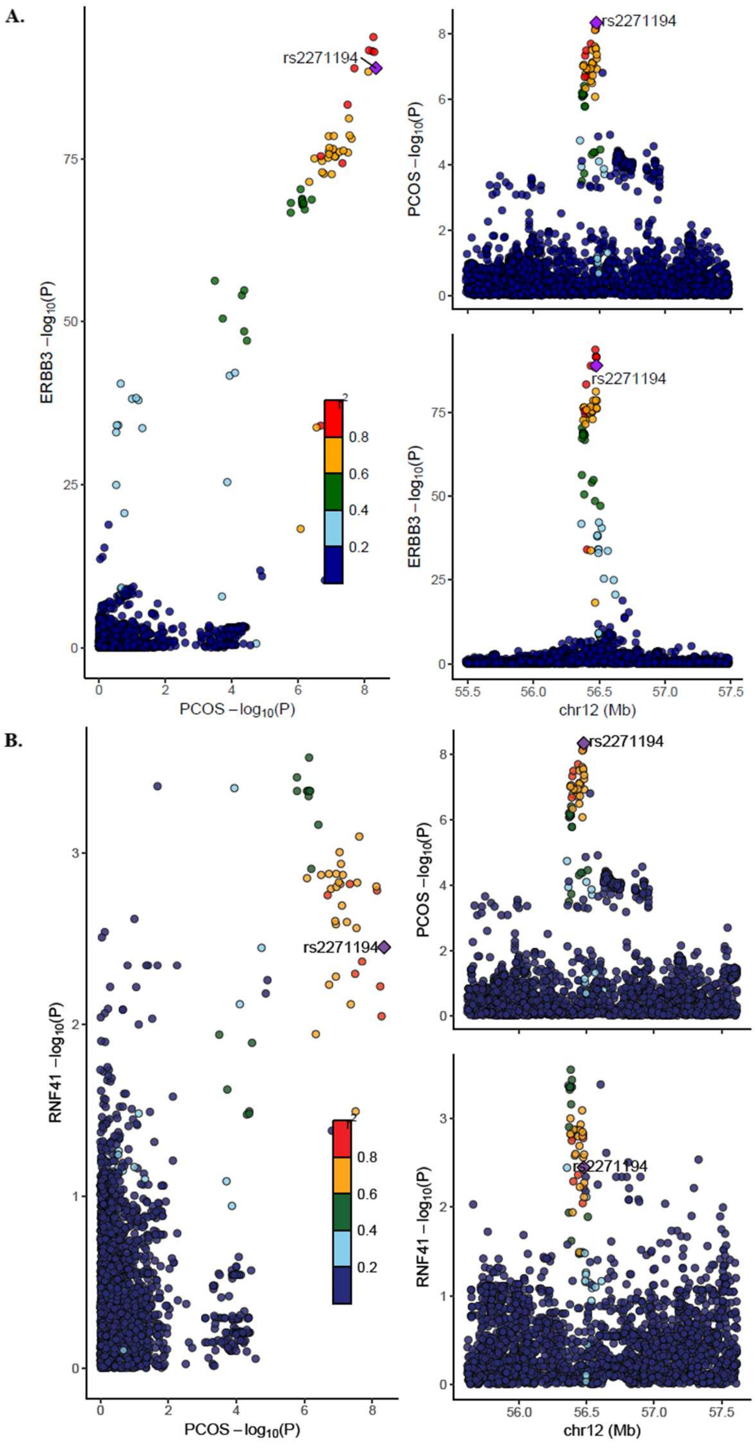


**Supplementary Figure 1A and 1B. Associations between genetic variants and PCOS risk, using Combined PCOS dataset, +/- 1 Mb region sizes for (A) *ERBB3* expression levels in blood (eQTLgen) (B) *RNF41* expression levels in coronary artery (GTEx)**

In each plot, each dot is a genetic variant. The SNP with the most significant P-value for PCOS is marked, with the other SNPs colour-coded according to linkage disequilibrium ($r^2$) in Europeans with the lead variant. SNPs with missing linkage disequilibrium information are also coded dark blue. In the left panels, -log10 P-values for associations with PCOS risk are on the x-axes, and -log10 P-values for associations with the expression levels on the y-axes. On the right panels, genomic positions are on the x-axes, and the y-axes show -log10 P-values for PCOS on the upper panel and -log10 P-values with the expression levels on the lower panel for the corresponding region. PCOS, polycystic ovary syndrome; SNP, single nucleotide polymorphism.

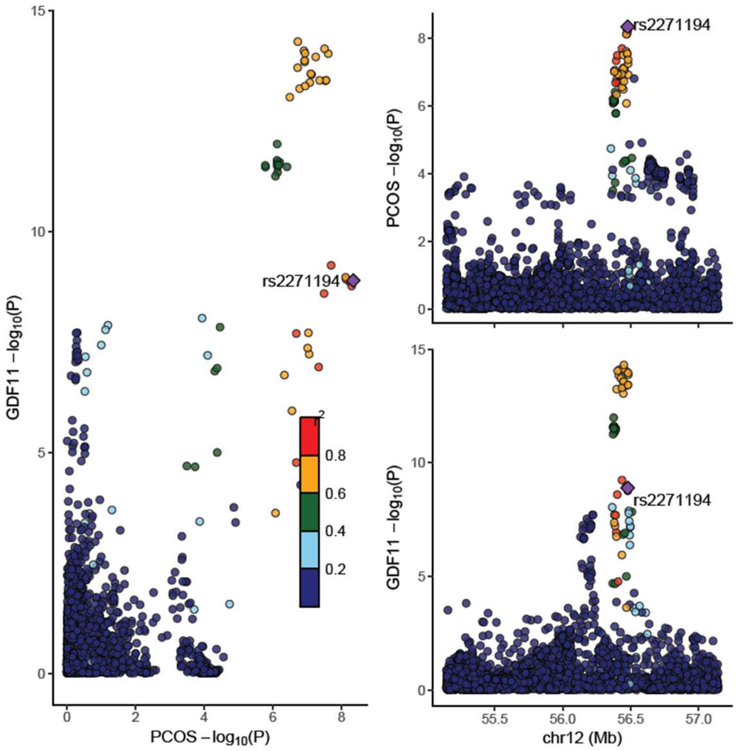


**Supplementary Figure 2. Associations between genetic variants and PCOS risk, using Combined PCOS dataset, +/- 1 Mb region sizes for *GDF11* expression levels in blood (eQTLgen)**
In each plot, each dot is a genetic variant. The SNP with the most significant P-value for PCOS is marked, with the other SNPs colour-coded according to linkage disequilibrium ($r^2$) in Europeans with the lead variant. SNPs with missing linkage disequilibrium information are also coded dark blue. In the left panel, -log10 P-values for associations with PCOS risk are on the x-axis, and -log10 P-values for associations with the expression levels on the y-axes. On the right panels, genomic positions are on the x-axes, and the y-axes show -log10 P-values for PCOS on the upper panel and -log10 P-values with the expression levels on the lower panel for the corresponding region. PCOS, polycystic ovary syndrome; SNP, single nucleotide polymorphism.

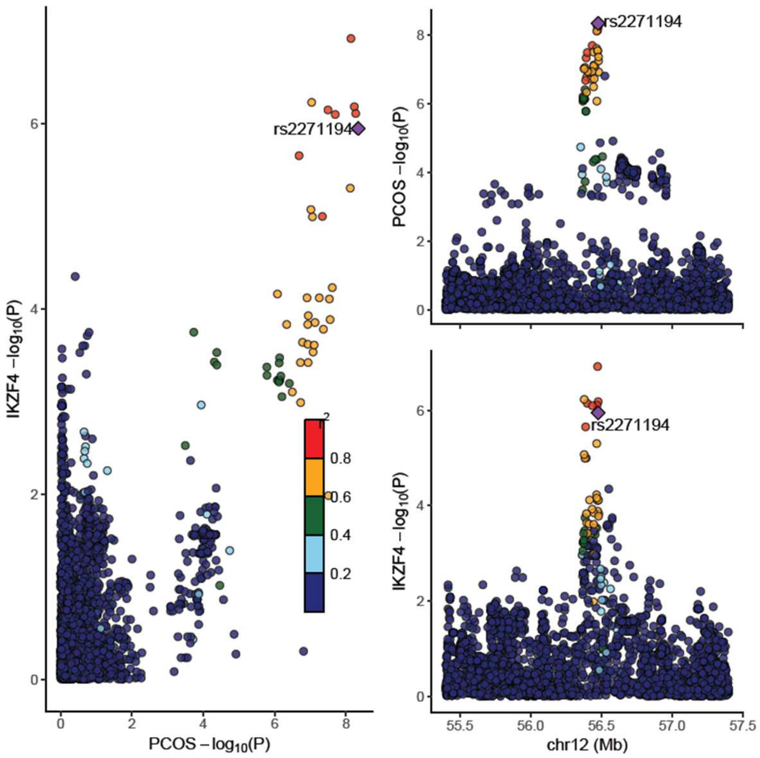


**Supplementary Figure 3. Associations between genetic variants and PCOS risk, using Combined PCOS dataset, +/- 1 Mb region sizes for *IKZF4* expression levels in esophagus (mucosa)**

In each plot, each dot is a genetic variant. The SNP with the most significant P-value for PCOS is marked, with the other SNPs colour-coded according to linkage disequilibrium ($r^2$) in Europeans with the lead variant. SNPs with missing linkage disequilibrium information are also coded dark blue. In the left panel, -log10 P-values for associations with PCOS risk are on the x-axis, and -log10 P-values for associations with the expression levels on the y-axes. On the right panels, genomic positions are on the x-axes, and the y-axes show -log10 P-values for PCOS on the upper panel and -log10 P-values with the expression levels on the lower panel for the corresponding region. PCOS, polycystic ovary syndrome; SNP, single nucleotide polymorphism.

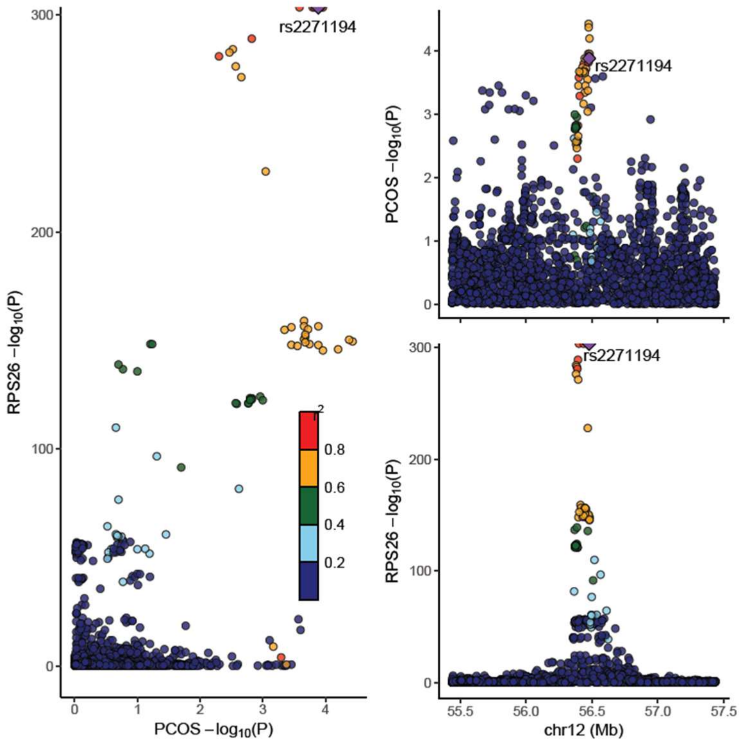


**Supplementary Figure 4. Associations between genetic variants and PCOS risk, using Combined PCOS dataset, +/- 1 Mb region sizes for *RPS26* expression levels in blood (eQTLgen)**
In each plot, each dot is a genetic variant. The SNP with the most significant P-value for PCOS is marked, with the other SNPs colour-coded according to linkage disequilibrium ($r^2$) in Europeans with the lead variant. SNPs with missing linkage disequilibrium information are also coded dark blue. In the left panel, -log10 P-values for associations with PCOS risk are on the x-axis, and -log10 P-values for associations with the expression levels on the y-axes. On the right panels, genomic positions are on the x-axes, and the y-axes show -log10 P-values for PCOS on the upper panel and -log10 P-values with the expression levels on the lower panel for the corresponding region. PCOS, polycystic ovary syndrome; SNP, single nucleotide polymorphism.

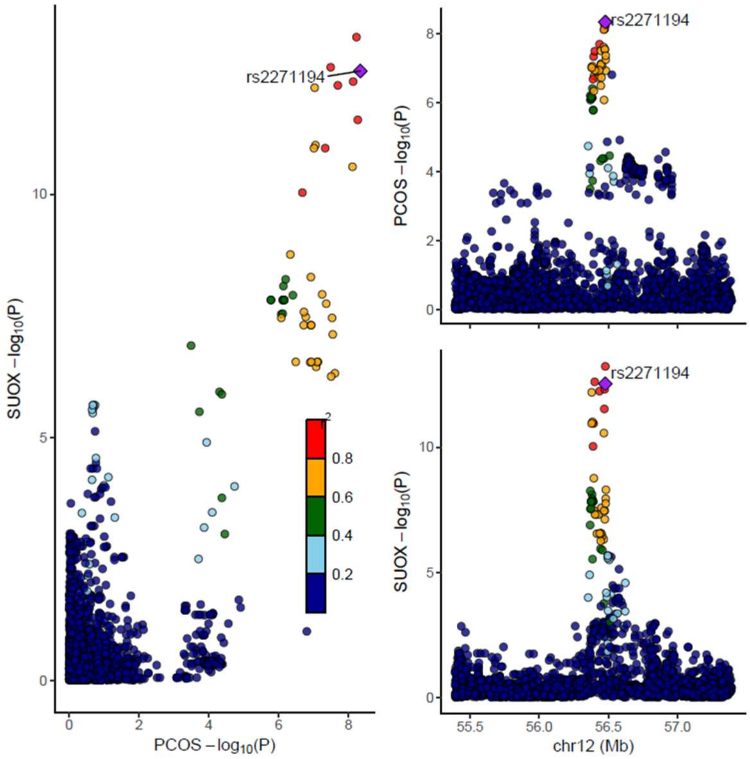


**Supplementary Figure 5. Associations between genetic variants and PCOS risk, using Combined PCOS dataset, +/- 1 Mb region sizes for *SUOX* expression levels in pituitary**

In each plot, each dot is a genetic variant. The SNP with the most significant P-value for PCOS is marked, with the other SNPs colour-coded according to linkage disequilibrium ($r^2$) in Europeans with the lead variant. SNPs with missing linkage disequilibrium information are also coded dark blue. In the left panel, -log10 P-values for associations with PCOS risk are on the x-axis, and -log10 P-values for associations with the expression levels on the y-axes. On the right panels, genomic positions are on the x-axes, and the y-axes show -log10 P-values for PCOS on the upper panel and -log10 P-values with the expression levels on the lower panel for the corresponding region. PCOS, polycystic ovary syndrome; SNP, single nucleotide polymorphism.

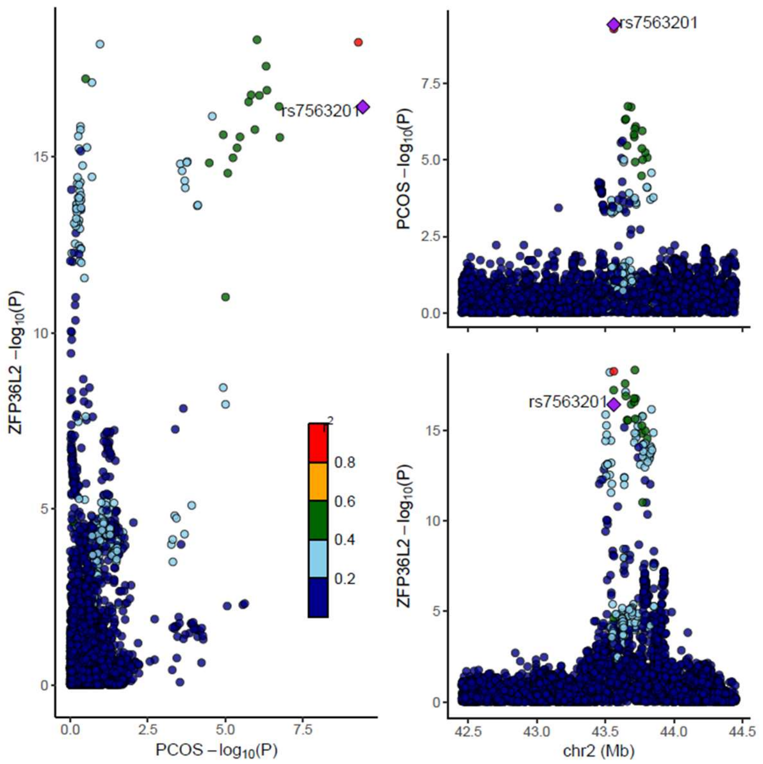


**Supplementary Figure 6. Associations between genetic variants and PCOS risk, using Combined PCOS dataset, +/- 1 Mb region sizes for *ZFP36L2* expression levels in blood (eQTLgen)**

In each plot, each dot is a genetic variant. The SNP with the most significant P-value for PCOS is marked, with the other SNPs colour-coded according to linkage disequilibrium ($r^2$) in Europeans with the lead variant. SNPs with missing linkage disequilibrium information are also coded dark blue. In the left panel, -log10 P-values for associations with PCOS risk are on the x-axis, and -log10 P-values for associations with the expression levels on the y-axes. On the right panels, genomic positions are on the x-axes, and the y-axes show -log10 P-values for PCOS on the upper panel and -log10 P-values with the expression levels on the lower panel for the corresponding region. PCOS, polycystic ovary syndrome; SNP, single nucleotide polymorphism.

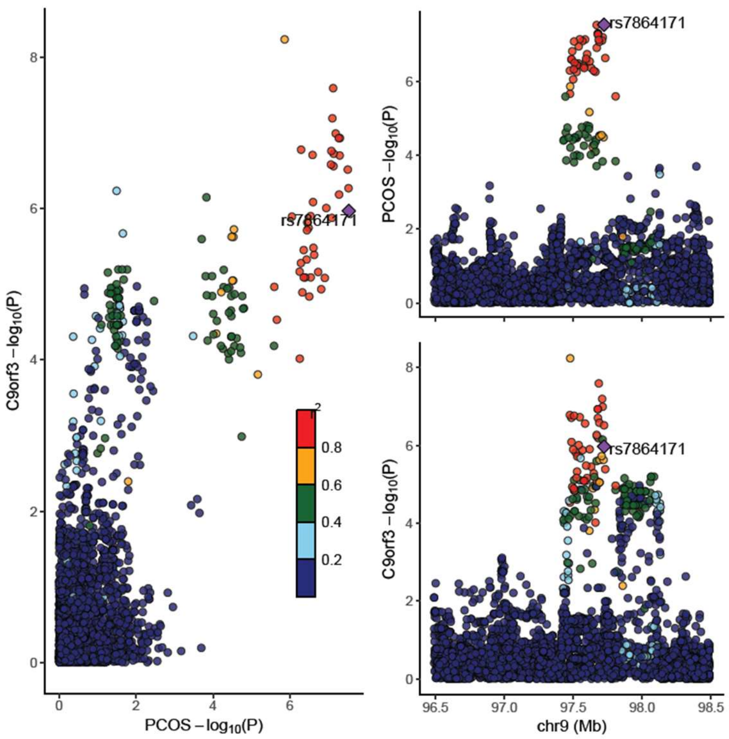


**Supplementary Figure 7. Associations between genetic variants and PCOS risk, using Combined PCOS dataset, +/- 1 Mb region sizes for *C9orf3* expression levels in atrial appendage of the heart**

In each plot, each dot is a genetic variant. The SNP with the most significant P-value for PCOS is marked, with the other SNPs colour-coded according to linkage disequilibrium ($r^2$) in Europeans with the lead variant. SNPs with missing linkage disequilibrium information are also coded dark blue. In the left panel, -log10 P-values for associations with PCOS risk are on the x-axis, and -log10 P-values for associations with the expression levels on the y-axes. On the right panels, genomic positions are on the x-axes, and the y-axes show -log10 P-values for PCOS on the upper panel and -log10 P-values with the expression levels on the lower panel for the corresponding region. PCOS, polycystic ovary syndrome; SNP, single nucleotide polymorphism.

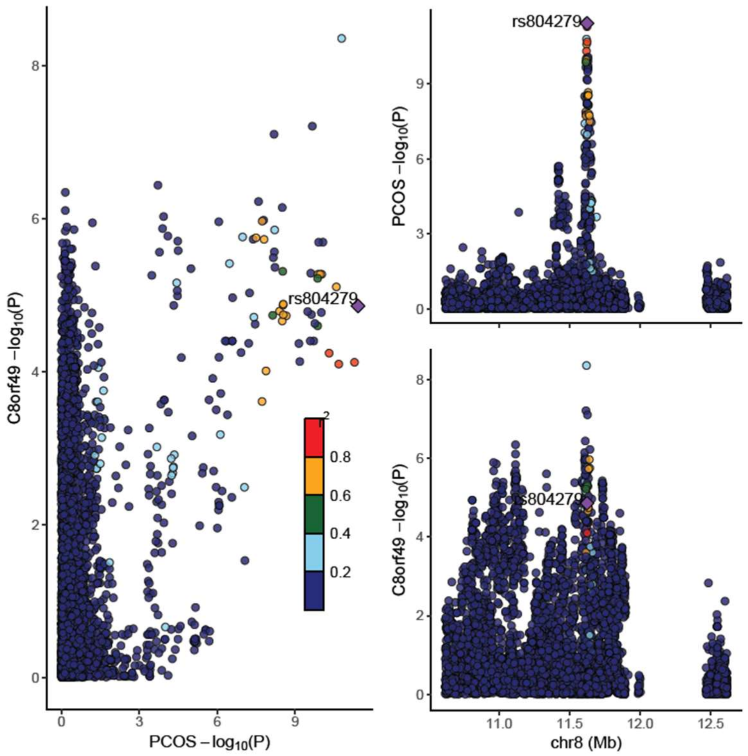


**Supplementary Figure 8. Associations between genetic variants and PCOS risk, using Combined PCOS dataset, +/- 1 Mb region sizes for *C8orf49* expression levels in stomach**
In each plot, each dot is a genetic variant. The SNP with the most significant P-value for PCOS is marked, with the other SNPs colour-coded according to linkage disequilibrium ($r^2$) in Europeans with the lead variant. SNPs with missing linkage disequilibrium information are also coded dark blue. In the left panel, -log10 P-values for associations with PCOS risk are on the x-axis, and -log10 P-values for associations with the expression levels on the y-axes. On the right panels, genomic positions are on the x-axes, and the y-axes show -log10 P-values for PCOS on the upper panel and -log10 P-values with the expression levels on the lower panel for the corresponding region. The region included an area without SNPs, spanning around 12-12.5 Mb on chromosome 8. A look-up using the UCSC genome browser (40) showed that the region had very poor mappability, explaining the lack of data. PCOS, polycystic ovary syndrome; SNP, single nucleotide polymorphism.

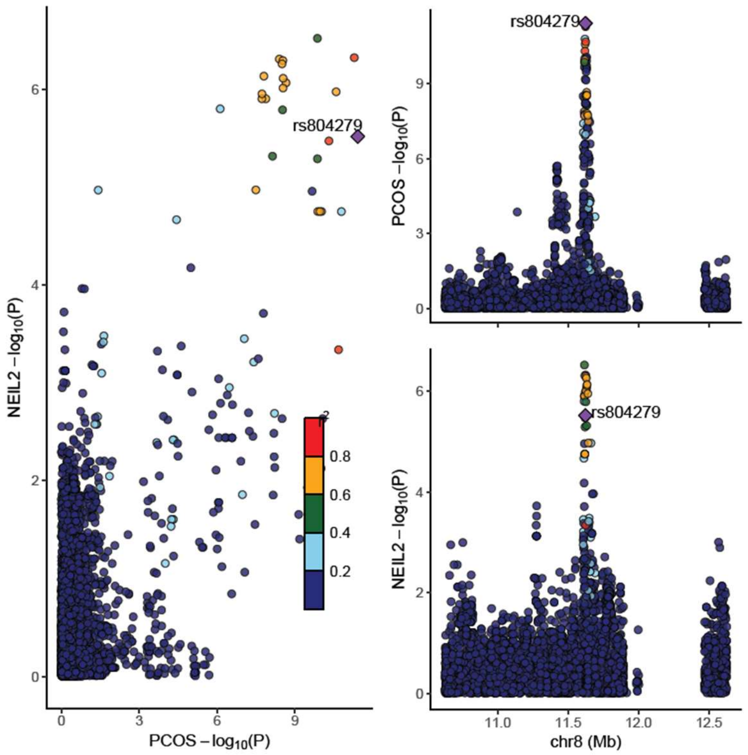


**Supplementary Figure 9. Associations between genetic variants and PCOS risk, using Combined PCOS dataset, +/- 1 Mb region sizes for *NEIL2* expression levels in EBV-transformed lymphocytes**

In each plot, each dot is a genetic variant. The SNP with the most significant P-value for PCOS is marked, with the other SNPs colour-coded according to linkage disequilibrium ($r^2$) in Europeans with the lead variant. SNPs with missing linkage disequilibrium information are also coded dark blue. In the left panel, -log10 P-values for associations with PCOS risk are on the x-axis, and -log10 P-values for associations with the expression levels on the y-axes. On the right panels, genomic positions are on the x-axes, and the y-axes show -log10 P-values for PCOS on the upper panel and -log10 P-values with the expression levels on the lower panel for the corresponding region. The region included an area without SNPs, spanning around 12-12.5 Mb on chromosome 8. A look-up using the UCSC genome browser (40) showed that the region had very poor mappability, explaining the lack of data. PCOS, polycystic ovary syndrome; SNP, single nucleotide polymorphism.

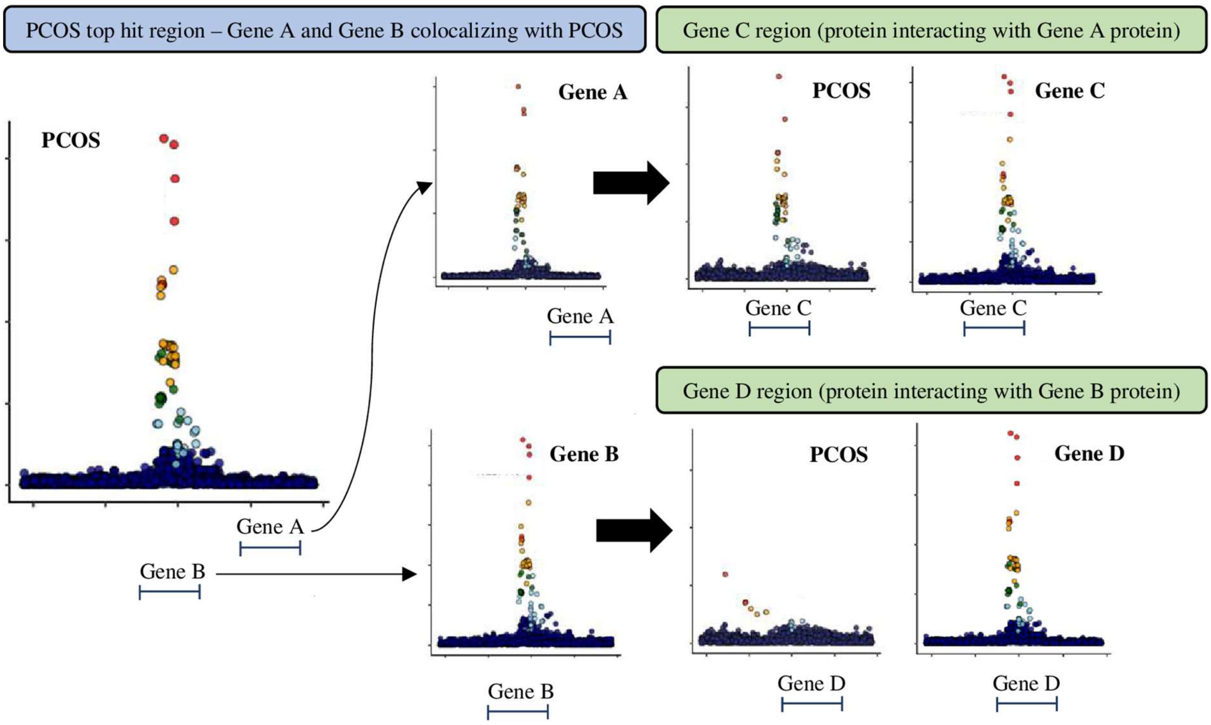


**Supplementary Figure 10. Rationale for investigating the evidence for colocalization between PCOS and the protein/expression levels of other genes interacting with the originally identified genes.**

For a region associated with PCOS P<5×10$^{-8}$, quantitative trait loci (QTLs) of Gene A and Gene B are both colocalizing with PCOS risk, possibly due to shared regulatory mechanisms. To investigate the evidence for each gene, the evidence for colocalization between PCOS and gene expression/protein levels for proteins with evidence of interaction with proteins of Gene A (denoted Gene C) and Gene B (denoted Gene D) are evaluated. If the QTL for Gene C colocalizes with PCOS (even though the top PCOS signal in the region of Gene C has a P-value >5×10$^{-8}$), whereas there is no colocalization between the QTL for Gene D and PCOS, we reasoned that this would provide more evidence for Gene A rather than Gene B being implicated in PCOS pathophysiology. PCOS, polycystic ovary syndrome; QTL, quantitative trait locus.

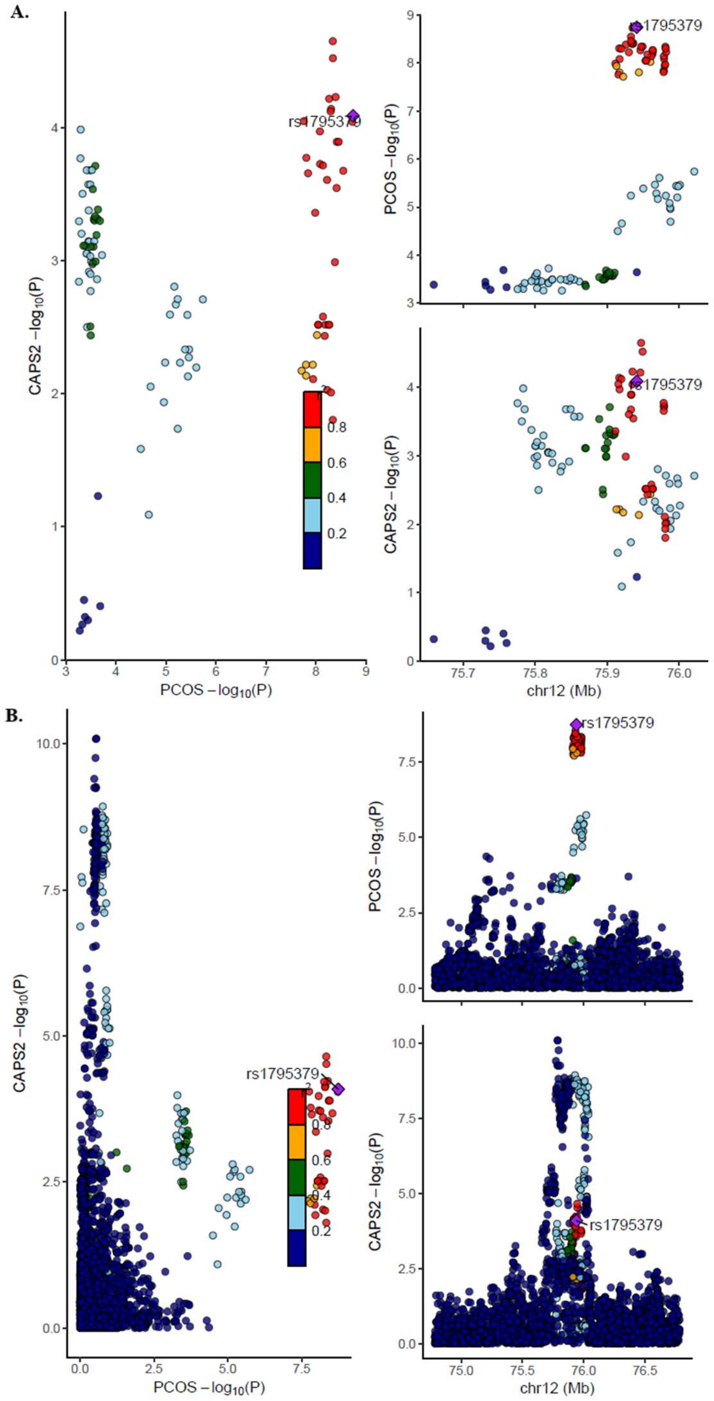


**Supplementary Figure 11A and 11B. Associations between genetic variants and PCOS risk, using Combined PCOS dataset for *CAPS2* expression levels in transverse colon using (A) Top 10,000 SNPs PCOS dataset (B) Combined PCOS dataset**

In each plot, each dot is a genetic variant. The SNP with the most significant P-value for PCOS is marked, with the other SNPs colour-coded according to linkage disequilibrium ($r^2$) in Europeans with the lead variant. SNPs with missing linkage disequilibrium information are also coded dark blue. In the left panels, -log10 P-values for associations with PCOS risk are on the x-axes, and -log10 P-values for associations with the protein/transcript levels on the y-axes. On the right panels, genomic positions are on the x-axes, and the y-axes show -log10 P-values for PCOS on the upper panel and -log10 P-values with the protein/expression levels on the lower panel for the corresponding region. PCOS, polycystic ovary syndrome; SNP, single nucleotide polymorphism.

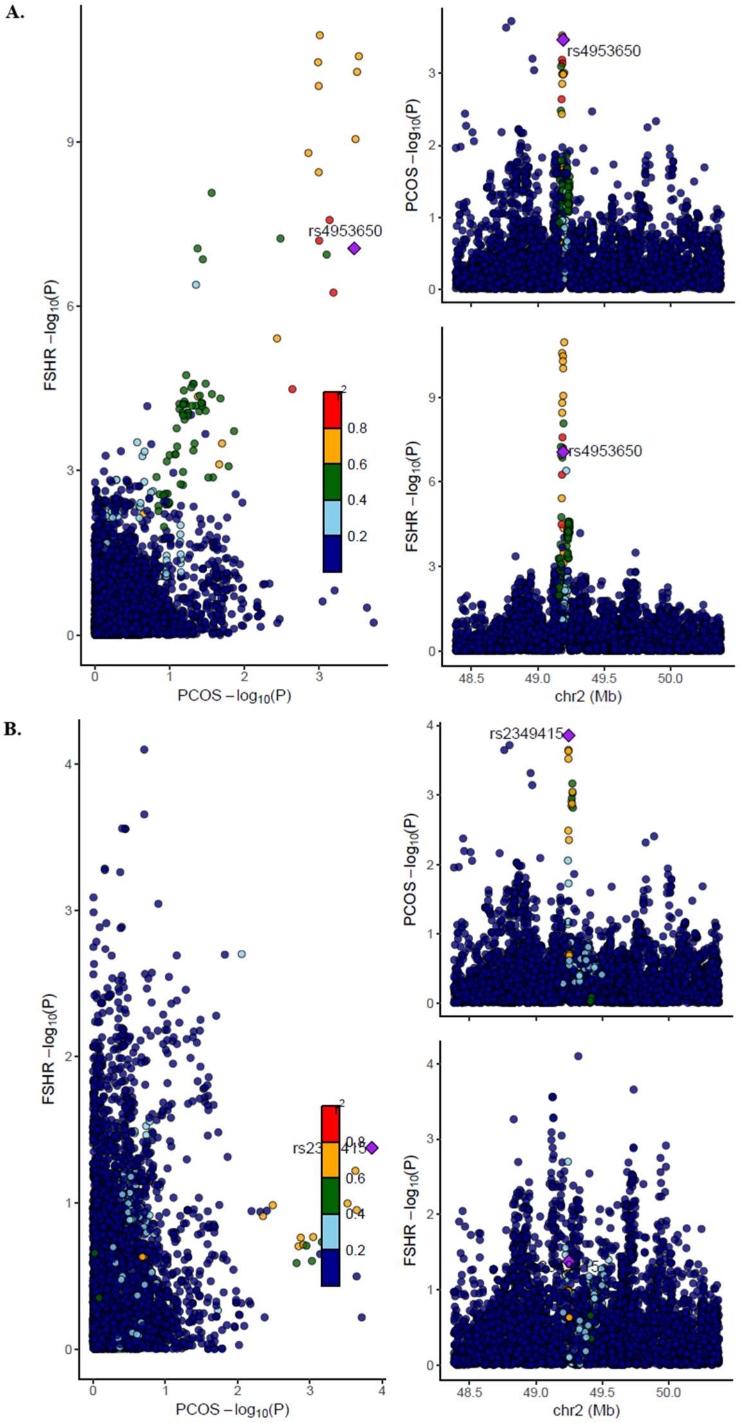


**Supplementary Figure 12A and 12B. Associations between genetic variants and PCOS risk, using Combined PCOS dataset, +/- 1 Mb region sizes for (A) *FSHR* expression levels in testis using estimates conditioned on rs2349415 (B) *FSHR* expression levels in testis using estimates conditioned on rs4953650**

In each plot, each dot is a genetic variant. The SNP with the most significant P-value for PCOS in the unconditioned Combined datasets is marked, with the other SNPs colour-coded according to linkage disequilibrium ($r^2$) in Europeans with the lead variant. SNPs with missing linkage disequilibrium information are also coded dark blue. In the left panels, -log10 P-values for associations with PCOS risk are on the x-axes, and -log10 P-values for associations with the transcript levels on the y-axes. On the right panels, genomic positions are on the x-axes, and the y-axes show -log10 P-values for PCOS on the upper panel and -log10 P-values with the expression levels on the lower panel for the corresponding region. FSH, follicle stimulating hormone; PCOS, polycystic ovary syndrome; SNP, single nucleotide polymorphism.

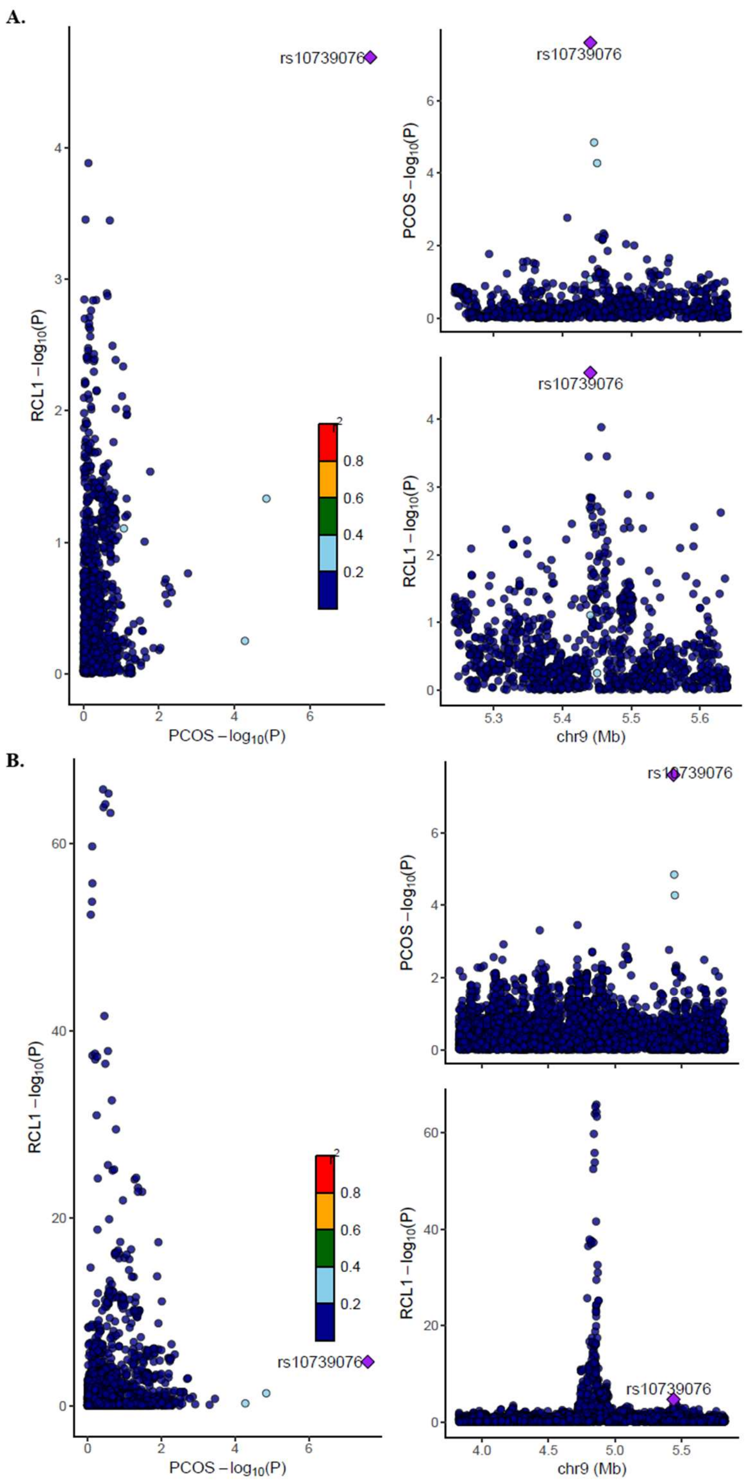


**Supplementary Figure 13A and 13B. Associations between genetic variants and PCOS risk, using Combined PCOS dataset for *RCL1* expression levels in blood (eQTLgen) using (A) +/- 200 kb region size (B) +/- 1 Mb region size**

In each plot, each dot is a genetic variant. The SNP with the most significant P-value for PCOS is marked, with the other SNPs colour-coded according to linkage disequilibrium ($r^2$) in Europeans with the lead variant. SNPs with missing linkage disequilibrium information are also coded dark blue. In the left panels, -log10 P-values for associations with PCOS risk are on the x-axes, and -log10 P-values for associations with the protein/transcript levels on the y-axes. On the right panels, genomic positions are on the x-axes, and the y-axes show -log10 P-values for PCOS on the upper panel and -log10 P-values with the protein/expression levels on the lower panel for the corresponding region. PCOS, polycystic ovary syndrome; SNP, single nucleotide polymorphism.
